# Improved and Simplified Diagnosis of Covid-19 using TE Extraction from Dry Swabs

**DOI:** 10.1101/2020.05.31.126342

**Authors:** Uday Kiran, C. G. Gokulan, Santosh Kumar Kuncha, Dhiviya Vedagiri, Karthik Bharadwaj Tallapaka, Rakesh K Mishra, Krishnan Harinivas Harshan

## Abstract

Rigorous testing is the way forward to fight the Covid-19 pandemic. Here we show that the currently used and most reliable RT-PCR based SARS-CoV-2 procedure can be further simplified to make it faster, safer and economical by bypassing the RNA isolation step. The modified method is not only fast and convenient but also at par with the traditional method in terms of accuracy, and therefore, can be used for mass screening. Our method takes about half the time and is cheaper by about 40% compared to current most widely used method. We also provide a variant of the new method that increases the efficiency of detection by about 20% compared to the currently used method. Taken together, we demonstrate a more effective and reliable method of SARS-CoV-2 detection.

## INTRODUCTION

Rapidly growing number of coronavirus disease 2019 (COVID-19) cases warrants reliable and quicker testing methods (Wu et al., 2020). Currently, Reverse Transcription-Polymerase Chain Reaction (RT-PCR) is the standard method being used for SARS-CoV-2 detection, largely owing to its high sensitivity (Al-Tawfiq and Memish, 2020; Emery et al., 2004). In the absence of specific drug and/or vaccine, the only way to control SARS-CoV-2 spread is large scale screening and isolation of the affected individuals at early stage of infection. Screening using antibody-based methods is rapid but cannot be used for early stage detection (Carter et al., 2020). Despite being a superior method, RT-PCR demands significant amount of time due to a laborious and expensive RNA isolation step. Currently, the challenge is to adapt a detection method which is quicker and still retaining the sensitivity of the standard RT-PCR-based method.

Here we show that the RNA isolation step for performing RT-PCR can be completely bypassed by extracting biological samples from dry swabs using TE buffer, which is cost-effective and can be used as a quick screening procedure. In addition, we also show that the sensitivity of the entire RT-PCR based detection is enhanced by at least 20%, by using RNA isolated from TE buffer extract compared to the traditional method.

## RESULTS AND DISCUSSION

We first hypothesized that SARS-CoV-2 nucleic acid could be detected directly by using viral transport medium (VTM) containing swabs of COVID-19 patients. This methodology involves the lysis of the virions (in VTM) by heating a 50 µl aliquot of VTM at 98°C for 6 minutes, followed by using 4 µl of this as a template for subsequent RT-PCR reaction.However, our results showed a reduced detection efficiency at 50% (n=24) compared to the traditional RNA isolation-based method (Supplementary Figure 1).

Although our data puts forth the feasibility of using VTM instead of extracted RNA, the detection ability of this method is limited to samples with moderate to high viral load. One of the probable reasons for this decreased efficiency could be dilution of the samples in 3 ml VTM, and to overcome this, we changed our sample collection strategy. To test the new strategy, two nasopharyngeal swab samples were collected from each of the fourteen patients with one swab transported dry and the other in VTM. The samples were processed as mentioned in the methods section (Figure 1A). Of the fourteen patients, five were tested negative and the remaining nine were positive samples, which were used for comparison. The results revealed that the performance of dry swab-TE buffer extracts in direct RT-PCR was at par with the currently used standard method of detection which has the additional RNA extraction step from VTM samples (Supplementary Table 1 – D1-D14).

**Figure 1:**
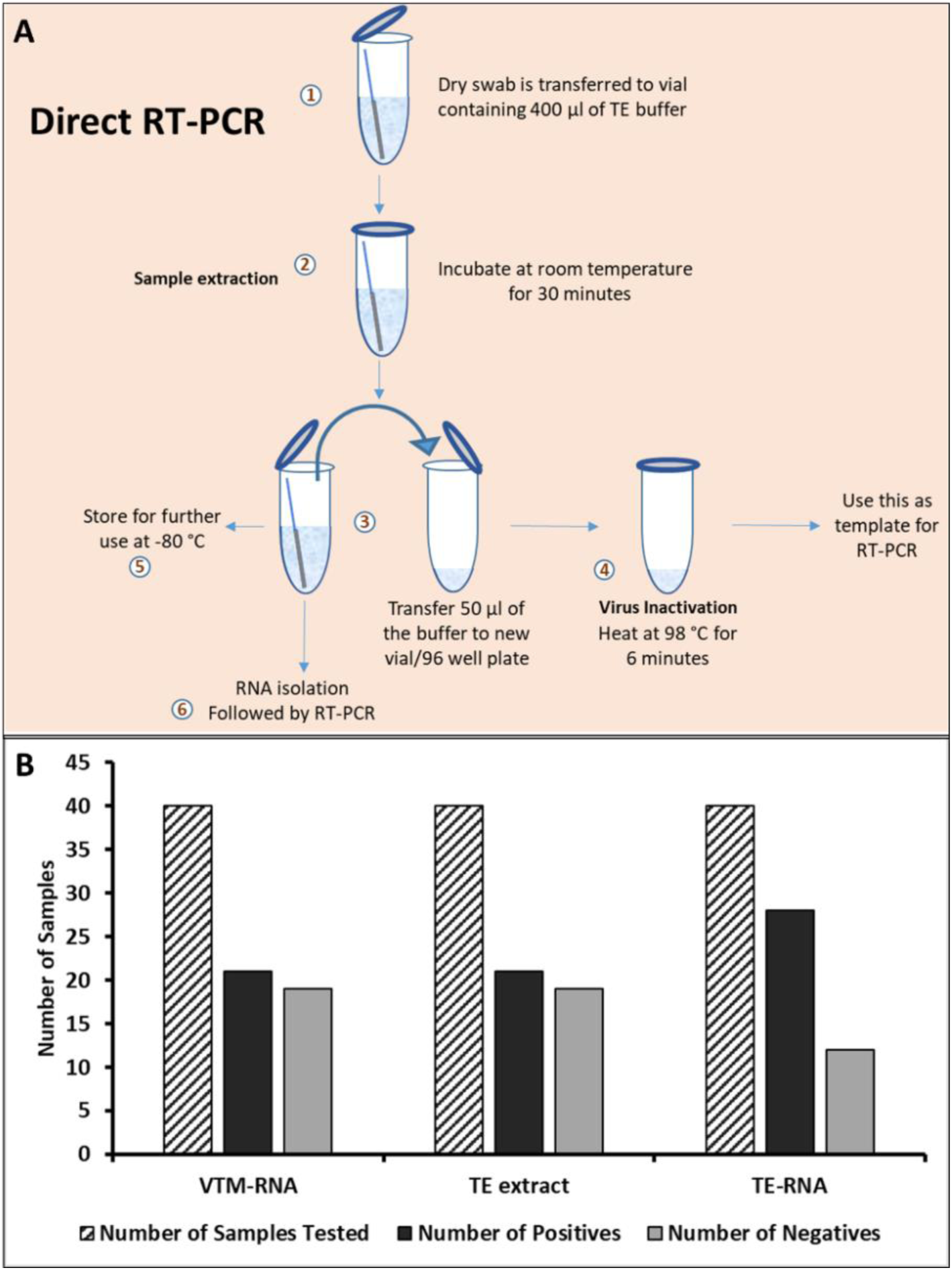
RNA extracted from TE buffer outperform other methods. A) Schematic of the entire protocol for TE-based sample extraction and RT-PCR. B) Bar graph representing the sensitivity of different methods for detecting the SARS-CoV-2 infection (n=40). Details in Supplementary Table 1.

To further validate the usage of TE buffer extract as a template for direct RT-PCR, we obtained similar samples from twenty-six patients, on the whole taking the sample size to forty. The results have further strengthened our observation that the TE buffer extract is as sensitive as the extracted RNA (n=40; Figure 1B). In fact, the average Ct values for TE-based RT-PCR were comparable to that of the traditional method and therefore can serve as an alternative method (Supplementary Figure 2). This approach can be employed as rapid and economical method for diagnosis which does not require any RNA extraction step. Our results are in line with the earlier reports of RNA extraction-free RT-PCR (Alcoba-Florez et al., 2020; Bruce et al., 2020; Smyrlaki et al., 2020), but here we have done a very extensive (n=40) and thorough standardization of this procedure which is now consistent and compelling.

One of the biggest challenges in diagnostics is overcoming the problem of false-negatives, and SARS-CoV-2 is not an exception to this. Recent reports have shown that the percentage of false negative reported for SARS-CoV-2 is between 20% and 40% with the onset of symptoms and varies with respect to the phase of infection (Kucirka et al., 2020; Li et al., 2020), which is alarming and calls for immediate improvements in the detection methodology. To address this issue, we have combined the TE-extraction method with traditional method that follows RNA-extraction. Here RNA was first isolated from TE buffer after extracting from the dry swabs (described in methodology), followed by RT-PCR. We were pleasantly surprised that almost one-third of the samples (6 of 19) which were consistently negative with traditional VTM-based method and also direct-RT-PCR method turned out to be positive for SARS-CoV-2. This observation was reproducible in multiple rounds of testing (Figure 1B, 2A). Upon a further closer look at the overall data, it was intriguing to note that the samples which were positive in the TE-based RNA extraction (and negative in other two methods) had a Ct value for only one of the two gene (E gene and RdRP), therefore possibly hinting at the low viral load which can be now picked by the new method. The increased detection limits are also evident in the decreased Ct values (3 Ct units on an average) (Figure 2B). To rule out any discrepancies in the sample processing we have used RNaseP as an internal control (Supplementary Table 2). Interestingly as an indication of RNA amount and quality, the RNaseP Ct values in case of TE-based approach had lower values compared to RNA isolated from VTM, thereby proving the higher efficiency of the TE-based approach (Supplementary Table 2). Therefore, the new hybrid method of TE-based sample extraction results in increasing the overall efficiency by at least 20% (Figure 2A; Supplementary Table 1). These results provide a remarkable improvement in the detection of SARS-CoV-2 patients with less viral load and therefore provides better opportunity to manage the pandemic.

**Figure 2:**
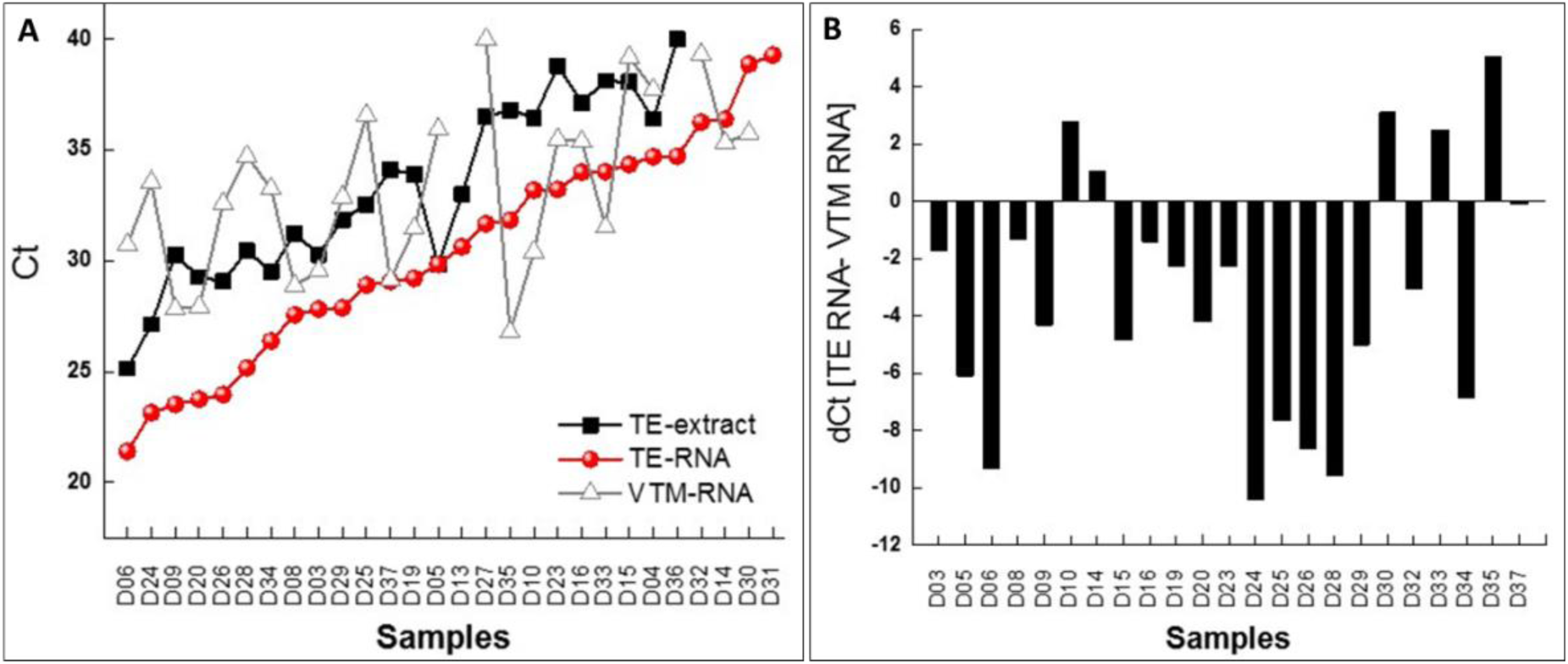
Increased sensitivity of COVID-19 diagnosis using TE-extracted RNA. A) Scatter plot of the Ct values for each sample obtained using different methods as labelled in the figure and the data points represents the average of Ct values of E and RdRP genes. B) Bar graph representing the difference in Ct values between TE-based RNA isolation compared to that of traditional (VTM-based RNA isolation) method. Negative and positive value indicates increased and reduced sensitivity, respectively, of the TE-based approach. The data points are average of two independent technical replicates.

Based on the above results, we recommend a 2-tier screening method for SARS-CoV-2 management. Since, TE buffer extracts can be used for direct RT-PCR without compromising the sensitivity of detection, we strongly recommend that this be employed as a first line of SARS-CoV-2 for large scale screening, while the TE buffer extract-based RNA could be employed if the former method yields an ambiguous result. This method can probably be expanded for screening other respiratory viral infections that are diagnosed using RT-PCR as well. Finally, we also recommend sample collection using dry-swab approach which not only eliminated the need of VTM, but also makes the sample handling, shipping and testing more convenient and safer for the frontline healthcare workers and technicians.

## MATERIALS AND METHODS

### Sample collection and transport

The swab samples were collected from voluntary patients at Gandhi Medical College & Hospital, Secunderabad, India. Two nasopharyngeal swabs were collected from each patient and one was transported as dry swab and another in VTM respectively and the samples were kept at 4° C till further processing.

### Sample processing

Complete sample processing was done in the BSL-3 facility of CSIR-CCMB by following Standard Operating Procedures.

a. Resuspension/extraction of biological material from dry swabs: The dry swabs were transferred to 1.5 ml microfuge tubes containing 400 μl of TE buffer [Tris pH-7.4 10 mM, EDTA 0.1 mM], The swabs were cut to make them fit into the tubes and incubated at room temperature for 30 min to ensure the release of biological material.
b. Heat Inactivation 50 µl of the sample was aliquoted from the VTM and TE buffer vials containing swabs and heated at 98° C for 6 min on dry heat block. The inactivated samples were directly used as a template for RT-PCR.

### RNA isolation

The RNA isolation from 3 ml VTM and TE-buffer (containing dry swab) was done by using the QIAamp Viral RNA isolation kit by following the manufacturer protocol. In both cases, 150 μl of the sample was processed for RNA-extraction.

### RT-PCR

All the RT-PCR work carried out in a BSL-2 facility of CSIR-CCMB, Hyderabad, India.

Heat inactivated VTM, TE buffer extract, and RNA isolated from TE buffer extract and VTM from respective samples were tested using the FDA approved LabGun COVID-19 detection RT-PCR kit (LabGenomics Co., Ltd., Republic of Korea). The primer-probe targets E and RdRP genes. The conditions for RT-PCR were followed according to the manufacturer protocol. All the reactions were multiplexed and an amount of 4 µl of the template was used per reaction. RT-PCR was performed in duplicates using LightCycler® 480 II (Roche Life Science, Germany) and the average values of both the experiments were used for the analysis. For plotting purposes Ct value mean of E and RdRP gene were used (Figure 2 and Supplementary Figure 2).

Excel/Origin software tool was used to generate all the plots/images in the manuscript.

An SOP for the entire protocol for implementation at the testing centres is provided as separate file “<u>Direct RT-PCR Method SOP</u>”.

## Supporting information

SOP for the RT-PCR method

## Ethical Statement

The study follows the institutional ethics committee guidelines.

## Acknowledgements

We acknowledge the state government official and the Department of Medical Education, Telangana for providing the samples for this study. UK, CGG and SKK thank the financial support received from CSIR, UGC and DST (INSPIRE), India respectively. All the authors acknowledge the support received from CSIR, India.

---

**Supplementary Figure 1:**
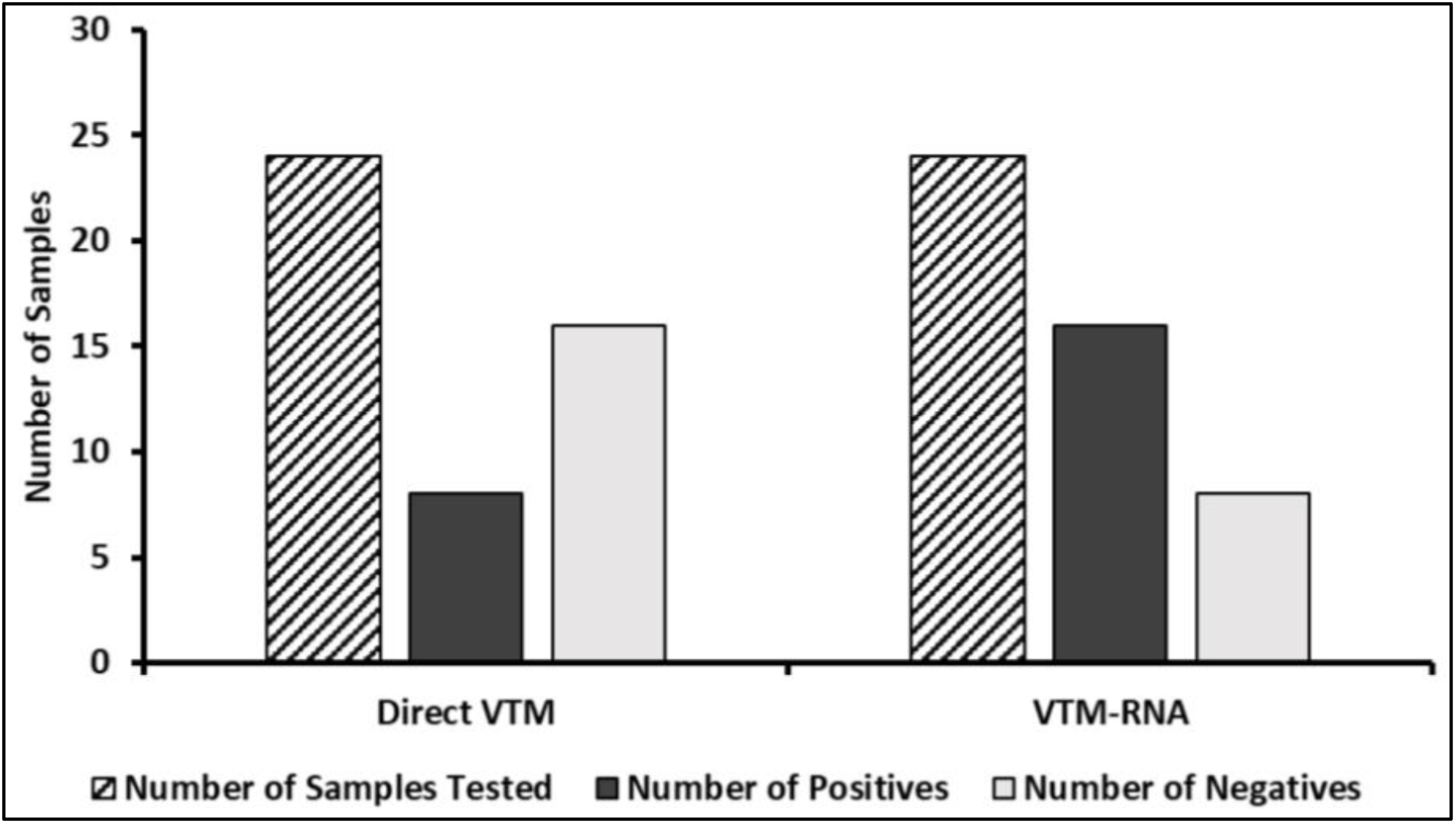
Direct RT-PCR from VTM is less sensitive comparable to traditional method. Bar graph showing the number of positives and negatives obtained using direct RT-PCR using VTM and RT-PCR performed using isolated RNA (n=24).

**Supplementary Figure 2:**
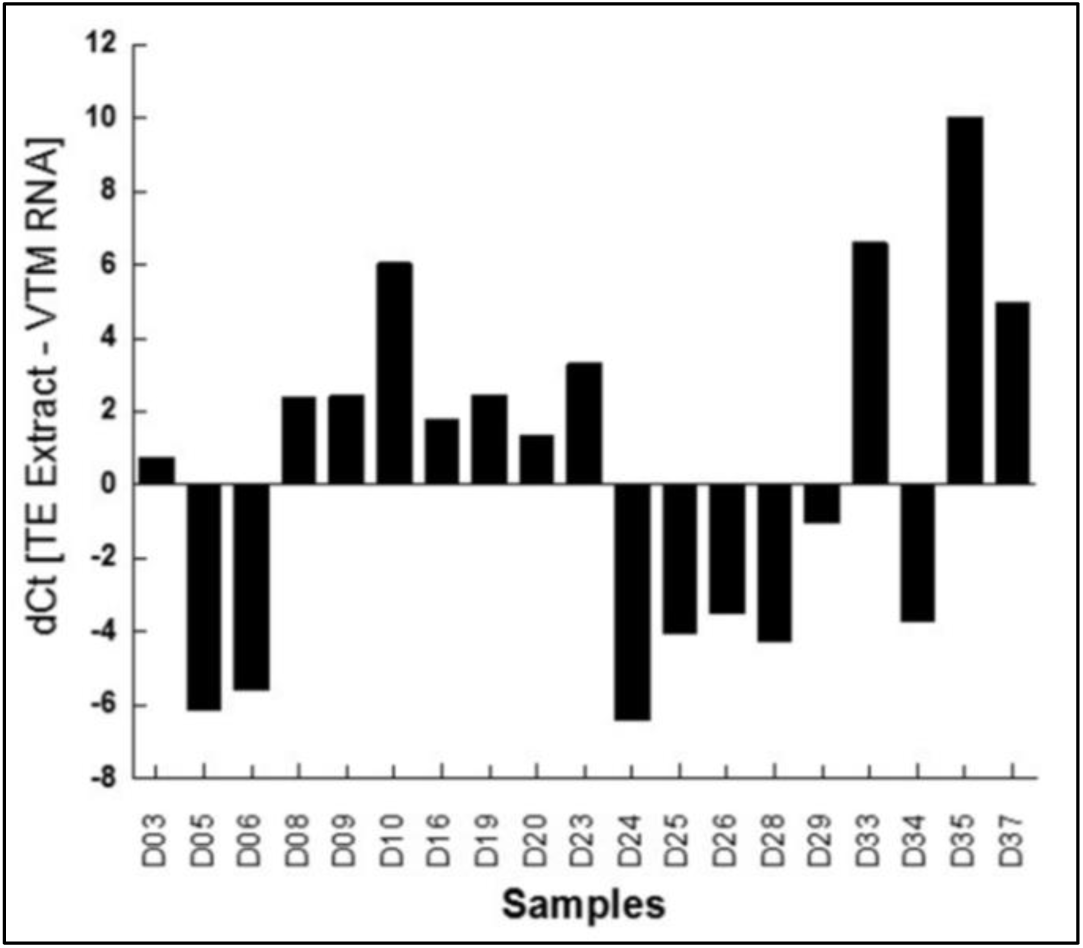
RNA extracted from TE buffer is comparable to the conventional based. Bar graph representing the difference in Ct values between TE-based direct RT-PCR and traditional VTM-based RNA isolation method. The negative values indicate higher efficiency of the technique while positive indicates lower sensitivity.

**Supplementary Table 1:** TE-based RNA isolation is the best approach.

| Samples | E (Average) |  |  | RdRP (Average) |  |  | Interpretation |  |  |
| --- | --- | --- | --- | --- | --- | --- | --- | --- | --- |
|  | TE Heated | TE RNA | VTM RNA | TE Heated | TE RNA | VTM RNA | TE Heated | TE RNA | VTM RNA |
| D01 |  |  |  |  |  |  | NEG | NEG | NEG |
| D02 |  |  |  |  |  |  | NEG | NEG | NEG |
| D03 | 29.93 | 27.74 | 28.91 | 30.63 | 27.9 | 30.19 | POS | POS | POS |
| D04 | 34.82 | 33.96 |  | 37.99 | 35.425 | 37.73 | POS | POS | NEG |
| D05 | 28.73 | 29.75 | 36.03 | 30.915 | 29.93 | 35.865 | POS | POS | POS |
| D06 | 24.83 | 21.3 | 30.53 | 25.435 | 21.495 | 30.92 | POS | POS | POS |
| D07 |  |  |  | 30.54 |  |  | NEG | NEG | NEG |
| D08 | 30.81 | 27.28 | 28.85 | 31.65 | 27.85 | 28.9 | POS | POS | POS |
| D09 | 29.77 | 23.55 | 27.51 | 30.75 | 23.49 | 28.155 | POS | POS | POS |
| D10 | 36.30 | 33.19 | 30.17 | 36.59 | 33.19 | 30.66 | POS | POS | POS |
| D11 |  |  |  |  |  |  | NEG | NEG | NEG |
| D12 |  |  |  |  |  |  | NEG | NEG | NEG |
| D13 | 32.94 | 30.63 |  | 33.08 | 30.63 |  | POS | POS | NEG |
| D14 |  | 36.39 | 35.24 |  | 36.39 | 35.41 | NEG | POS | POS |
| D15 | 38.06 | 34.37 | 38.36 |  | 34.32 | 40.00 | NEG | POS | POS |
| D16 | 37.55 | 33.68 | 35.59 | 36.75 | 34.31 | 35.19 | POS | POS | POS |
| D17 |  |  |  |  | 34.47 |  | NEG | NEG | NEG |
| D18 |  | 37.67 |  |  |  |  | NEG | NEG | NEG |
| D19 | 34.09 | 29.29 | 31.40 | 33.71 | 29.14 | 31.52 | POS | POS | POS |
| D20 | 29.33 | 23.76 | 27.79 | 29.20 | 23.74 | 28.08 | POS | POS | POS |
| D21 |  |  |  |  |  |  | NEG | NEG | NEG |
| D22 |  |  |  |  |  |  | NEG | NEG | NEG |
| D23 | 38.44 | 33.18 | 34.26 | 39.15 | 33.27 | 36.72 | POS | POS | POS |
| D24 | 27.13 | 23.01 | 33.39 | 27.13 | 23.28 | 33.71 | POS | POS | POS |
| D25 | 32.64 | 28.88 | 36.05 | 32.43 | 28.94 | 37.10 | POS | POS | POS |
| D26 | 29.60 | 23.95 | 32.42 | 28.56 | 23.98 | 32.77 | POS | POS | POS |
| D27 | 35.98 | 31.78 |  | 37.04 | 31.56 | 40.00 | POS | POS | NEG |
| D28 | 30.59 | 25.12 | 34.73 | 30.31 | 25.21 | 34.72 | POS | POS | POS |
| D29 | 31.95 | 27.84 | 32.09 | 31.77 | 27.92 | 33.68 | POS | POS | POS |
| D30 |  | 37.73 | 35.13 |  | 40.00 | 36.38 | NEG | POS | POS |
| D31 |  | 38.91 |  |  | 39.65 |  | NEG | POS | NEG |
| D32 |  | 36.08 | 40.00 |  | 36.44 | 38.64 | NEG | POS | POS |
| D33 | 38.22 | 33.68 | 31.39 | 38.03 | 34.36 | 31.66 | POS | POS | POS |
| D34 | 29.41 | 26.19 | 32.95 | 29.63 | 26.58 | 33.56 | POS | POS | POS |
| D35 | 36.57 | 31.85 | 26.71 | 37.06 | 31.81 | 26.85 | POS | POS | POS |
| D36 | 40.00 | 34.81 |  |  | 34.63 |  | NEG | POS | NEG |
| D37 | 33.90 | 28.98 | 29.02 | 34.36 | 29.16 | 29.28 | POS | POS | POS |
| D38 |  |  |  |  |  |  | NEG | NEG | NEG |
| D39 |  |  |  |  |  |  | NEG | NEG | NEG |
| D40 |  |  |  |  |  |  | NEG | NEG | NEG |

List of all the Ct values obtained using different approaches.

**Supplementary Table 2:** RNaseP values indicating the integrity of RNA samples.

| Samples | TE Heated | TE-RNA | VTM-RNA |
| --- | --- | --- | --- |
| D01 | 34.99 | 32.39 | 34.75 |
| D02 | 32.34 | 30.7 | 33.43 |
| D03 | 31.43 | 29.21 | 31.49 |
| D04 | 27.79 | 24.66 | 30.97 |
| D05 | 29.56 | 28.46 | 33.09 |
| D06 | 30.16 | 28.37 | 33.05 |
| D07 | 34.4 | 32.11 | 36.7 |
| D08 | 29.87 | 29.27 | 32.78 |
| D09 | 31.8 | 30.53 | 32.87 |
| D10 | 31.89 | 29.82 | 34.80 |
| D11 | 32.25 | 30.99 | ND |
| D12 | 30.72 | 29.83 | 33.9 |
| D13 | 32.82 | 32.3 | 36.02 |
| D14 | 29.54 | 28.52 | 33.3 |
| D15 | 33.25 | 30.32 | 33.51 |
| D16 | 32.8 | 30.93 | 33.54 |
| D17 | 33.68 | 31.23 | 33.66 |
| D18 | 32.15 | 29.62 | 34.62 |
| D19 | 31.16 | 28.36 | 34.76 |
| D20 | 31.77 | 27.77 | 34.74 |
| D21 | 34.03 | 30.22 | 34.64 |
| D22 | 28.07 | 23.23 | 31.13 |
| D23 | 31.43 | 28.29 | 35.56 |
| D24 | 30.67 | 27.65 | 32.85 |
| D25 | 28.81 | 26.25 | 34.83 |
| D26 | 26.58 | 24.34 | 34.99 |
| D27 | 30.47 | 25.72 | 33.26 |
| D28 | 32.58 | 30.18 | 33.37 |
| D29 | 27.61 | 26.6 | 26.11 |
| D30 | 31.1 | 27.95 | 29.53 |
| D31 | 34.93 | 31.93 | 36.44 |
| D32 | 30.65 | 28.76 | 35.92 |
| D33 | 30.7 | 29.11 | 35.61 |
| D34 | 32.81 | 29.88 | 35 |
| D35 | 30.44 | 26.69 | 32.56 |
| D36 | 33.42 | 31.78 | 35.06 |
| D37 | 29.12 | 27.54 | 32.9 |
| D38 | 34.33 | 31.07 | 33.08 |
| D39 | 34.56 | 31.04 | 33.29 |
| D40 | 31.77 | 27.29 | 34.21 |

List of all the Ct values of RNaseP (internal control). ND: not detected

