## Supplementary material for "Improved and Simplified Diagnosis of Covid-19 using TE Extraction from Dry Swabs": SOP for the RT-PCR method

### SOP for Direct RT-PCR based detection of SARS-CoV2

#### *Background:*

Current method of Covid-19 testing is based on the initial extraction of RNA from virus transport medium (VTM) in which the nasopharyngeal/oropharyngeal swabs are placed, followed by Taqman RT-PCR to detect viral RNA genome. This is a robust process and by far the best method for detection of virus in persons. However, there are several limitations that restrict the extent of its utility. One of the major flaws is the initial sample handling at the place of testing where patient samples need to be catalogued and aliquoted for RNA extraction. At this stage, the personnel handling the samples are exposed to potentially SARS-CoV-2 positive VTMs and any incident of aerosolization of VTM can expose the personnel to virus and infection. Another major drawback is the stage of RNA extraction. In addition to being expensive and short supply, this is also the major rate limiting step since one person can prepare RNAs from about twenty-four samples in 2.30 hrs.

We present here an improved method sample collection and processing that makes the Covid-19 testing process safer, faster and cheaper.

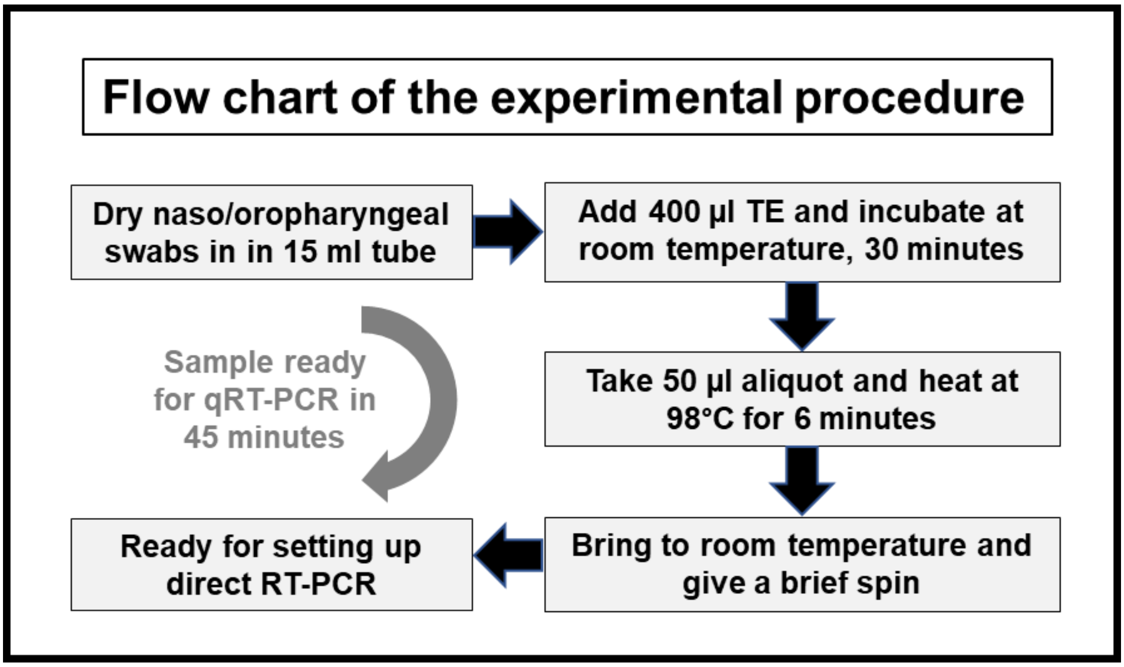

**Protocol for sample collection and processing for Direct RT-PCR**

| | | |
|---|---|---|
| 1 | Collect the naso/oropharyngeal swabs in empty 15 ml tubes. | |
| 2 | Pack the sample tubes in a single zip lock cover packing. There is no need to use parafilm wrap. | |
| 3 | Transport samples to the testing centre preferably on ice. | |
| 4 | The samples can be processed immediately or can be stored at 4°C for 24 hours. (freeze the samples if the test is not being conducted in next 2 days). | |
| 5 | Add 400 µl of TE buffer to the dry swab containing tube and incubate at room temperature for 30 minutes. Follow next step depending whether a 96 well plate of 1.5 ml tube is going to be used. | |
| | For 96 well plates | For 1.5 ml tubes |
| 6 | Transfer 50 µl of the sample to 96 well plate. | Transfer 50 µl of the sample to labelled 1.5 ml tube. |
| 7 | Seal the 96 well plate and heat it in PCR at 98 °C for 6 minutes and give a brief spin to the plate. | Heat the samples at 98°C for 6 minutes using a heating block and give a brief spin to the tube. |
| 8 | Use the samples directly for setting up the RT-PCR (as per the manufacturer guidelines). Please note all the steps needs to be carried out in hood. | |

**Requirements:**

*Recurring*: Swabs, 15 ml falcons (or equivalent) tubes, TE (10 mM Tris pH 7.4, 0.1 mM EDTA buffer prepared in nuclease free water), Common plasticware.

*Non-recurring:* Dry bath/heat block (for processing in 1.5 ml tubes), PCR machine (if processing using 96 well plates), Multichannel pipettes (for processing in 96 well plates)

**Comparison of traditional method with Direct RT-PCR method**

| **Major differences in the sample processing procedure** | | |
|---|---|---|
| | Traditional Method | Direct RT-PCR |
| Swabs collected | 3 ml VTM | Dry swabs |
| Packing | Multilayer packing individual samples to avoid leakage. | Multiple tubes can be packed into a single zip lock cover. |
| Unpacking | Very laborious.<br>High risk of infection due to liquid spillages (if any) | Simple.<br>No liquid spillage and hence low risk of infection |
| Inactivation of sample and lysis | Lysis takes longer time due to air bubbles | Extract the sample in TE |
| RNA isolation | Yes | No needed |
| Setting up of RT-PCR | Samples are independently added into the 96 well plate | Samples are added to the 96 well plate using multichannel pipette |

| **Time and Man Power requirement** (for 100 samples) | | |
|---|---|---|
| | Traditional Method | Direct RT-PCR |
| Unpacking & cataloguing | 3 personnel for 3 hours | 2 personnel for 1 hour |
| Inactivation & lysis | 2 personnel for 3 hours | 2 personnel for 3 ½ hours |
| RNA isolation | 4 personnel for 2 ½ hours | |
| RT-PCR | 2 personnel for 3 ½ hours | |
| **Total Time** | **10 hours** | **< 5 hours** |
| **Total Man power** | **11** | **4** |

| **Estimated cost reduction by Direct RT-PCR method** (for 100 samples, in INR) | |
|---|---|
| Swab collection (VTM cost) | 20,000 |
| RNA isolation | 40,000 |
| RT-PCR (by multiplexing in case of Labgun kit) | 60,000 |
| **Total** | **1,20,000** |
